# Phosphoproteomic analysis of mammalian infective *Trypanosoma brucei* subjected to heat shock suggests atypical mechanisms for thermotolerance

**DOI:** 10.1101/2020.02.07.938761

**Authors:** Cher P. Ooi, Corinna Benz, Michael D. Urbaniak

## Abstract

The symptoms of African sleeping sickness, caused by the parasite *Trypanosoma brucei*, can include periods of fever as high as 41 °C which triggers a heat shock response in the parasite. To capture events involved in sensing and responding to heat shock in the mammalian infective form we have conducted a SILAC-based quantitative proteomic and phosphoproteomic analysis of *T. brucei* cells treated at 41 °C for 1h. Our analysis identified 193 heat shock responsive phosphorylation sites with an average of 5-fold change in abundance, but only 20 heat shock responsive proteins with average of 1.5-fold change. These data indicate that protein abundance does not rapidly respond (≤1 h) to heat shock, and that the changes observed in phosphorylation site abundance are larger and more widespread. The heat shock responsive phosphorylation sites showed enrichment of RNA binding proteins with putative roles in heat shock response included P-body / stress granules and the eukaryotic translation initiation 4F complex. The ZC3H11-MKT1 complex, which stabilises mRNAs of thermotolerance proteins, appears to represent a key signal integration node in the heat shock response.

## Introduction

*Trypanosoma brucei* are extracellular eukaryotic parasites transmitted by the bite of an infected Tsetse fly which cause African sleeping sickness in humans and the related wasting disease Nagana in livestock and wild game. There are currently 70 million people at risk across Africa, with agricultural losses projected at $2.5 billion over the next two decades due to this disease [1, 2], African sleeping sickness is fatal if allowed to progress to a late stage of the disease and there are few treatment alternatives if resistance to front line therapeutics arise [3].

Symptoms of African sleeping sickness include periods of fever, which in some cases can be as high as 41 °C [4]. Moreover, the body temperature of domestic cattle can increase up to 2 °C from physiological norms over the course of a day [5, 6], and hosts in the wild, such as African gazelles, can achieve body temperatures of up to 43 °C during physical exertion [7]. How do these extracellular parasites adapt to periods when the temperature of the host is elevated beyond normal physiological conditions? Deviation from physiological norms in temperature can be devastating to eukaryotic organisms. Protein folding is affected and misfolded proteins accumulate in the cytoplasm. Eukaryotic cells respond to heat shock (HS) in two ways: they trigger a general arrest in protein translation to prevent further accumulation of misfolded proteins, and increase the expression of HS proteins (Hsps), which aid protein folding and degradation, despite the global block in protein translation. The eukaryotic HS response is conserved in *T. brucei* [8-10], but the mechanisms mediating this response appear to be divergent.

In mammalian cells, the HS translation arrest is triggered by the phosphorylation of the translation factor eIF2α which blocks translation initiation. This mechanism is not conserved across all eukaryotic organisms, as the HS-triggered translation arrest in yeast and fruit flies is not via eIF2α phosphorylation [8, 11, 12]. It is unknown if phosphorylation of eIF2α triggers the translation arrest in *T. brucei* during HS as previous mutational analysis have yielded negative results [8, 13]. Therefore, the translation arrest during HS in *T. brucei* could be triggered either by an unknown phosphorylation of eIF2α or via an eIF2α–independent mechanism.

Hsps bind to exposed hydrophobic regions on misfolded proteins to stabilise or promote degradation. Five major groups are conserved across all organisms (Hsp60s, Hsp70s, Hsp90s, Hsp100s and small Hsps), and some have defined functions in eukaryotes. In yeast and mammals, the canonical Hsp70-Hsp40-Hsp110 network functions to disaggregate misfolded proteins following HS and labels them for ubiquitination and degradation [14], and Hsp104 forms a hexameric ring which threads misfolded proteins through its central pore to resolve protein aggregates that form following HS [15–17]. In *T. brucei, Hsp70* and *Hsp83* mRNAs increase following HS [8, 9], and Hsp70 has been implicated to be essential for the HS response in *T. brucei* [18]. However, it is unknown if the functions of these Hsps are conserved in *T. brucei*. Moreover, small Hsps (sHsps) have only recently been identified within the *T. brucei* genome [19]. One sHsp has been demonstrated to have a role in HS survival in *Leishmania* [20], but it is unclear if sHsps play a significant role in the HS response of *T. brucei*.

Up-regulation of Hsp expression is key to survival during HS, and this is regulated at the level of transcription in mammalian cells. HS increases binding of the transcription factor HSF1 to promoters upstream of *Hsp* genes, increasing the processivity of RNA polymerase II which up-regulates Hsp expression [21]. This mechanism of regulation cannot operate in *T. brucei* as genes with diverse functions are arranged in long polycistronic arrays under a common transcription promoter. Control of gene expression is primarily post-transcriptional, with RNA binding proteins (RBPs) regulating messenger RNA (mRNA) levels [22]. RBPs have been demonstrated to bind to some *Hsp* mRNAs in *T. brucei* with the likely role of stabilising them to maintain expression during HS. The zinc finger protein ZC3H11 is an RBP which has been extensively documented to stabilise HS responsive mRNAs in the insect infective (procyclic form, PCF) life cycle stage of *T. brucei* by binding to AU-rich elements [18, 23]. ZC3H11 is only essential in the mammalian infective (bloodstream form, BSF) life cycle stage of *T. brucei*, yet does not stabilise HS-related mRNAs to the same degree as in PCFs [23], suggesting that other mechanisms, such as other RBPs, may instead be involved. Moreover, Hsp regulation in kinetoplastids during HS may not rely primarily on increased levels of expression. A large proportion of phosphorylated proteins following HS in *Leishmania donovani* are Hsps and protein chaperones, including Hsp70, Hsp90 and the Hsp complex protein STI1. It has been suggested that the functions of these HS-associated proteins are phosphorylation-dependent, though this has only been experimentally demonstrated for STI1 [24–26].

In this work, we provide insight into the phosphoproteomic changes to BSF *T. brucei* during HS. Previous global-omics studies of HS in *T. brucei* have focused on its PCF life cycle stage [10, 27]. This work, to our knowledge, constitutes the first documentation of proteomic and phosphoproteomic changes in mammalian-infective BSF *T. brucei in* response to HS.

## Material and methods

### Trypanosome strains and culturing

All experiments were carried out with BSF *Trypanosoma brucei brucei* 427. Cells were cultured in HMI-9 medium supplemented with 15 % foetal calf serum. To assess cell viability after HS, cultures at a density of 10^5^ cells/ml were subjected to 41 °C for different amounts of time in a water bath before being transferred back to normal growth conditions at 37 °C. Cell viability was assessed by counting only motile cells with a Neubauer haemocytometer.

### RNA manipulation and qPCR analysis

Total RNA was isolated from cells treated to HS with a Qiagen RNeasy Mini Kit (Qiagen). Genomic DNA was removed using Turbo DNase (Ambion) according to manufacturer’s instructions. Synthesis of cDNA was carried out using an Omniscript RT kit (Qiagen) with 100 ng total RNA per reaction. qPCR reactions were assembled using Brilliant II SYBR low ROX master mix (Agilent) using 1 μl of diluted cDNA (1/100). Primers used are listed in supplementary Table S1. qPCR data are presented as fold changes using the Livak method [28].

### Stable isotope labelling of T. brucei cells

The stable isotope labelling by amino acids in cell culture (SILAC) labelling of BSF *T. brucei* cells was performed as described previously [29, 30]. In brief, log phase were diluted 10,000-fold into HMI11-SILAC (HMI9 medium lacking Serum plus, l-arginine and l-lysine) supplemented with 10% dialysed FCS (1000 MWCO, Dundee Cell Products) and the standard concentration of either normal isotopic abundance l-arginine and l-lysine (HMI11+ R0K0, referred to as Light) or ^13^C_6_,^15^N_4_ l-arginine and ^13^C_6_,^15^N_2_ l-lysine (HMI11 + R10K8, referred to as Heavy). The stable isotope-labelled amino acids were obtained from CK Isotopes, UK. After three days growth (>9 cell divisions), the cell reached ~2 × 10^6^ cells/ml, and the Light labelled cells were heat shocked at 41 °C for 1h in a water bath. Cells were harvested by centrifugation and hypotonically lysed at 1 × 10^9^ cells/mL for 5 min on ice in the presence 0.1 μM 1-chloro-3-tosylamido-7-amino-2-heptone (TLCK), 1 mM benzamidine, 1 mM phenyl-methyl sulfonyl fluoride (PMSF), 1 μg/mL leupeptin, 1 μg/mL aprotinin and Phosphatase Inhibitor Mixture II (Calbiochem), and stored at −80 °C prior to subsequent processing.

### Proteomic sample preparation

Samples were thawed and equal cell numbers (2 × 10^8^ cells per sample) combined, then the proteins solubilized with SDS and tryptic peptides generated by an adaptation of the filter aided sample preparation procedure as described previously [30, 31]. The digested peptide solution was then removed from the column, diluted to 3 ml with ABC and acidified with 0.1% TFA before desalting using a 500 mg C_18_ column (SepPak, Waters), lyophilisation and storage at −80 °C.

Separation of peptides from phosphopeptides using Fe-IMAC was performed as described by Ruprecht *et al* [32] with minor modifications as described previously [33]. Samples were subjected to HPLC (LC Packing Famos) on a Fe-IMAC column (4 × 50 mm ProPac IMAC-10, Thermo Fisher Scientific) with peptide eluted in Buffer A (30% MeCN, 0.07% TFA) and phosphopeptide eluted in a gradient of Buffer B (30% MeCN, 0.5% NH_4_OH) with elution monitored by absorbance at 280 nm. Early eluting fractions (<6 min) corresponding to peptides and later eluting fractions corresponding to phosphopeptides (~30 min) were lyophilised and stored at −80 °C prior to further processing.

Peptide and phosphopeptide were separately fractionated using a High pH Fractionation kit (ThermoFisher) according to the manufacturer’s instructions with the following modifications. Lyophilised (phospho)peptides were resuspended in 100μl 5 mM NH_4_OH and applied to the pretreated columns (MeCN, 0.1 % TFA, and two 5 mM NH_4_OH washes). The flow-through was reapplied before being collected and desalted using C_18_ microspin columns (Harvard Apparatus). The columns were then washed 3 times with 5 mM NH_4_OH and the (phospho)peptides eluted with different concentrations of MeCN (2, 3, 4, 6, 10 and 50% MeCN). The eluates were concatenated into five fractions as follows: F1 = 2% + 50% eluates, F2 = 3% eluate, F3 = 4% eluate, F4 = 6% eluate, F5 = 10% eluate + desalted flow through. The concatenated fractions were then lyophilised prior to analysis by LC-MS/MS.

### Mass Spectrometry Data Acquisition

Liquid chromatography tandem mass spectrometry (LC-MS/MS) was performed by the FingerPrints Proteomic Facility at the University of Dundee. Liquid chromatography was performed on a fully automated Ultimate U3000 Nano LC System (Dionex) fitted with a 1 × 5 mm PepMap C_18_ trap column and a 75 μm × 15 cm reverse phase PepMap C_18_ nanocolumn (LC Packings, Dionex). Samples were loaded in 0.1% formic acid (buffer A) and separated using a binary gradient consisting of buffer A (0.1% formic acid) and buffer B (90% MeCN, 0.08% formic acid). Peptides were eluted with a linear gradient from 5 to 40% buffer B over 65 min. The HPLC system was coupled to an LTQ Orbitrap Velos Elite mass spectrometer (Thermo Scientific) equipped with a Proxeon nanospray ion source. For phosphoproteomic analysis, the mass spectrometer was operated in data dependent mode to perform a survey scan over a range 335 – 1800 m/z in the Orbitrap analyzer (*R* = 60,000), with each MS scan triggering fifteen MS^2^ acquisitions of the fifteen most intense ions using multistage activation on the neutral loss of 98 and 49 Thomsons in the LTQ ion trap [34]. For proteomic analysis, the mass spectrometer was operated in data dependent mode with each MS scan triggering fifteen MS^2^ acquisitions of the fifteen most intense ions in the LTQ ion trap. The Orbitrap mass analyzer was internally calibrated on the fly using the lock mass of polydimethylcyclosiloxane at *m/z* 445.120025.

### Mass Spectrometry Data Processing

Data was processed using MaxQuant [35] version 1.6.1.0 which incorporates the Andromeda search engine [36]. Proteins were identified by searching a protein sequence database containing *T. brucei brucei* 927 annotated proteins (Version 37, 11,074 protein sequences, TriTrypDB [37], http://www.tritrypdb.org/) supplemented with frequently observed contaminants (porcine trypsin, bovine serum albumins and mammalian keratins). Search parameters specified an MS tolerance of 6 ppm, an MS/MS tolerance at 0.5 Da and full trypsin specificity, allowing for up to two missed cleavages. Carbamidomethylation of cysteine was set as a fixed modification and oxidation of methionine, *N*-terminal protein acetylation and *N*-pyroglutamate were allowed as variable modifications. The experimental design included matching between runs for the concatenated fractions in each experiment. Analysis of the phosphoproteomic data used the same parameters, except for the addition of phosphorylation of serine, threonine and tyrosine residues as variable modifications. Peptides were required to be at least 7 amino acids in length, with false discovery rates (FDRs) of 0.01 calculated at the levels of peptides, proteins and modification sites based on the number of hits against the reversed sequence database.

SILAC ratios for phosphorylation sites were calculated using only data from the phosphoproteomic experiments, and SILAC ratios for proteins were calculated using only data from the proteomic experiments. SILAC ratios were calculated where at least one peptide could be uniquely mapped to a given protein group and required a minimum of two SILAC pairs. To account for any errors in the counting of the number of cells in each sample prior to mixing, the distribution of SILAC ratios was normalised within MaxQuant at the peptide level so that the median of log_2_ ratios is zero, as described by Cox *et al*.[35]. We have deposited the Thermo RAW files and search results in ProteomeXchange (http://www.proteomexchange.org) consortium via the Pride partner repository [38] with the dataset identifier PXD016482, enabling researchers to access the data presented here.

### Statistical analyses

Statistical analyses of proteomics data was carried out using Perseus 1.6.1.3 [39] using data obtained from the function genomic database TriTrypDB [37]. Species that changed significantly upon HS were identified using intensity-weighted significance (Significance B) [35] with a Benjamini-Hochberg corrected FDR ≤ 0.01. Categorical enrichment was determined using Fisher’s exact test with a Benjamini-Hochberg corrected FDR ≤ 0.01. All other statistical analyses were carried out using GraphPad Prism 7.

## Results

### A Heat shock response is triggered in BSF T. brucei after incubation at 41 °C

To simulate HS, BSF *T. brucei* were incubated at 41 °C for up to 4 h, and cell viability immediately after heat treatment was determined by counting the cells that remained motile (Figure 1A.). Although incubation for 1 h at 41 °C leads to a slight decrease in live cells (22 %, p = 0.49), the number of live cells only decreased significantly after 2 h at 41 °C (74 %, p = 0.04) and no motile cells were observed after 4 h at 41 °C. To assess the recovery of cells from heat treatment, cells were returned to culture at 37 °C and their growth monitored for 72 h. As no motile cells were observable after 4 h of incubation at 41 °C, this treatment was omitted from this analysis. Cells incubated for 1 h at 41 °C remain viable after HS, albeit with a reduced growth rate up to 24 h post-treatment, and cells treated for 2 h at 41 °C continued to lose viability and failed to grow for up to 48 h posttreatment (Figure 1B.).

**Figure 1.**
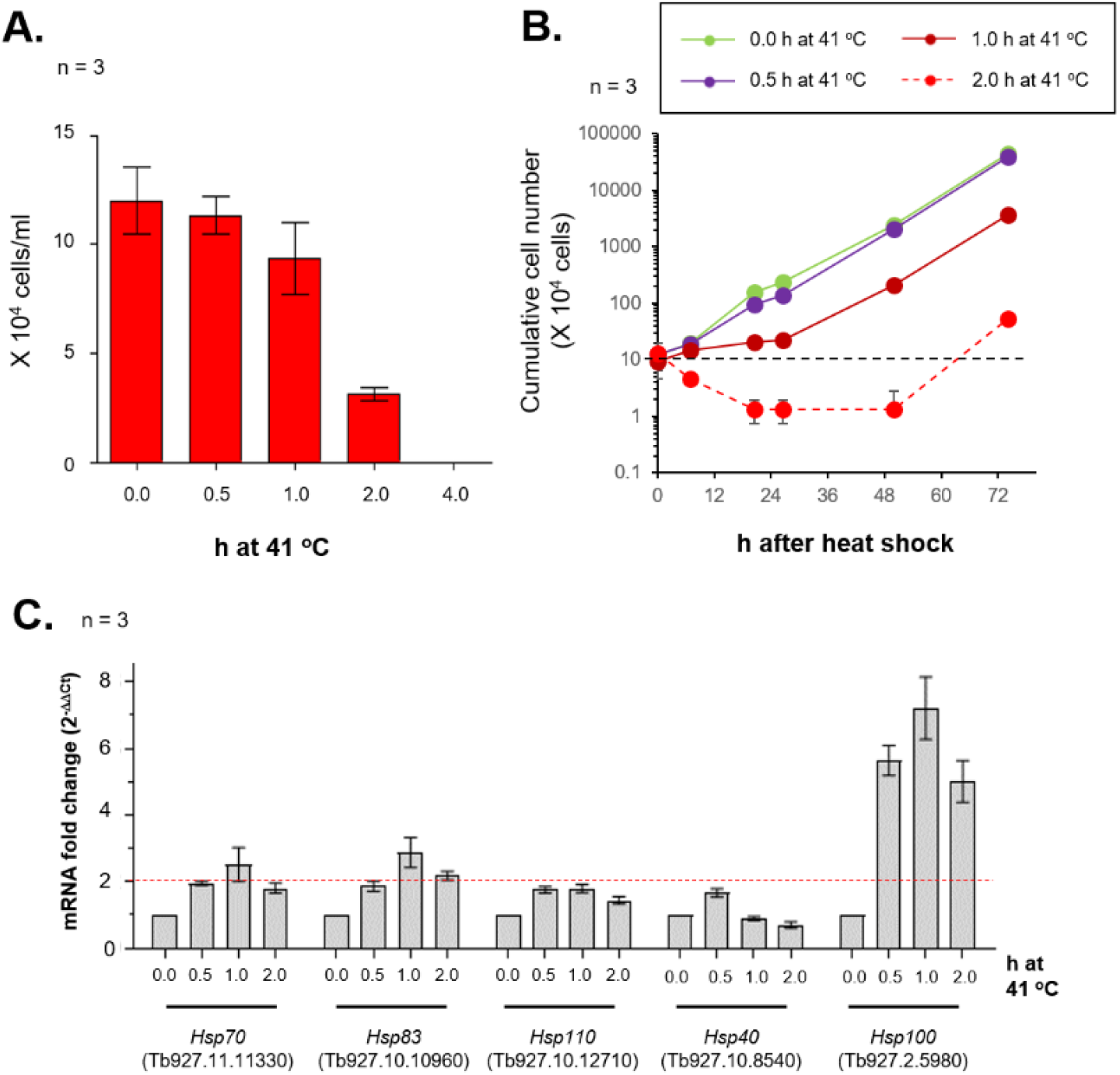
A heat shock response is triggered in BSF *T. brucei* after 1 h at 41 °C. **A.** Count of motile BSF *T. brucei* cells after heat shock (41 °C) for 0.0, 0.5, 1.0, 2.0 or 4.0 h. B. Cumulative growth curves of BSF *T. brucei* cells after 0.0, 0.5, 1.0, 2.0 or 4.0 h of heat shock. Cells were resuspended at 10^5^ cells/ml after heat shock and cultured in HMI9 at 37 °C. **C.** qPCR analysis of selected *Hsps* after heat shock. Data are presented as fold change (2^-ddCt^) compared to untreated cells (0. h at 41 °C). Error bars are of standard deviation from three biological replicates.

To determine if a HS response was triggered in these cells, total RNA was harvested from heat-treated cells for qPCR quantitation of *Hsp* mRNA. Total RNA quality was assessed by electrophoresis on a 2 % agarose gel, and only samples collected from cells incubated for 2 h at 41 °C showed signs of degradation (Figure S1A.). All qPCR quantitation used the 28S □ subunit of ribosomal RNA as the reference and the critical threshold (Ct) value for this transcript remained constant with all HS treatments (Figure S1B.). Five *Hsps* were selected for qPCR quantitation. These included two *Hsps* known to be up-regulated at the transcript level after HS in *T. brucei* [8, 9] (*Hsp70*, Tb927.11.11330 and *Hsp83*, Tb927.10.10960), homologues to other members of the canonical Hsp70-Hsp40-Hsp110 network in yeast and mammals [14] (*Hsp110*, Tb927.10.12710 and *Hsp40*, Tb927.10.8540), and the closest *T. brucei* homologue to yeast Hsp104, which is also up-regulated after HS in PCF *T. brucei* [15–18] (*Hsp100*, Tb927.2.5980). *Hsps* with a ≥ 2-fold increase were defined as up-regulated after heat shock. Consistent with previous reports [8, 9], *Hsp70* and *Hsp83* transcript levels increase > 2-fold after 1 h of HS (Figure 1C.), indicating that a HS response is triggered in the treated cells. Likewise, *Hsp100* transcript levels increase ≥ 6-fold after HS, suggesting that the *Hsp104* homologue has a role in the *T. brucei* HS response. The transcript level for other member of the Hsp70-Hsp40-Hsp110 network remain relatively unchanged, which may indicate a divergence of the *T. brucei* HS response from that of its mammalian host.

### Quantitative phosphoproteomic and proteomic analysis of heat shock sensing in BSF T. brucei

To dissect the HS response in BSF *T. brucei* in an unbiased manner, we conducted a SILAC-based quantitative proteomic and phosphoproteomic analysis of the BSF *T. brucei* heat shock response. We chose to examine an early time point (41 °C for 1h) to capture signal transduction events involved in sensing and responding to heat shock. Cells were grown in HMI11-SILAC supplemented with 10% dialysed FCS and either normal isotopic abundance l-arginine and l-lysine (Light) or ^13^C_6_,^15^N_4_ l-arginine and ^13^C_6_,^15^N_2_ l-lysine (Heavy) for >9 cell divisions to ensure steady state labelling, and then the Light cells were heat shocked at 41 °C for 1h. Heavy and Light cells were independently hypotonically lysed in the presence of phosphatase and protease inhibitors, then equal cell numbers (~2 × 10^8^) combined and subjected to FASP and tryptic digest [30, 31]. Phosphopeptides were separated from peptides using Fe-IMAC [32], and the two pools separately fractionated using the high pH reverse phase with concatenation prior to LC-MS/MS analysis [33].

Analysis of the phosphoproteomic data identified a total of 8,055 phosphorylation sites (localisation probability ≥ 0.75) on 2,381 proteins, and was able to quantify the change in abundance between the HS and untreated cells for 7,258 phosphorylation sites on 2,137 proteins (Supplementary Table S2). A total of 193 of phosphorylation sites on 148 proteins changed abundance significantly upon HS (FDR ≤ 0.01), with 138 sites increasing in abundance with an average 6-fold increase and 55 sites decreasing in abundance by an average of 4-fold (Figure 2A). Proteomic analysis identified a total of 4,002 proteins, and was able to quantify the change in abundance of 2,581 proteins between the HS and untreated cells (Supplementary Table S3). The overall changes in protein abundance were modest, with only 20 proteins showing significant changes (FDR < 0.01), with 7 proteins increasing in abundance by an average of 1.4-fold and 13 proteins decrease in abundance by an average of 1.5-fold (Figure 2B).

**Figure 2.**
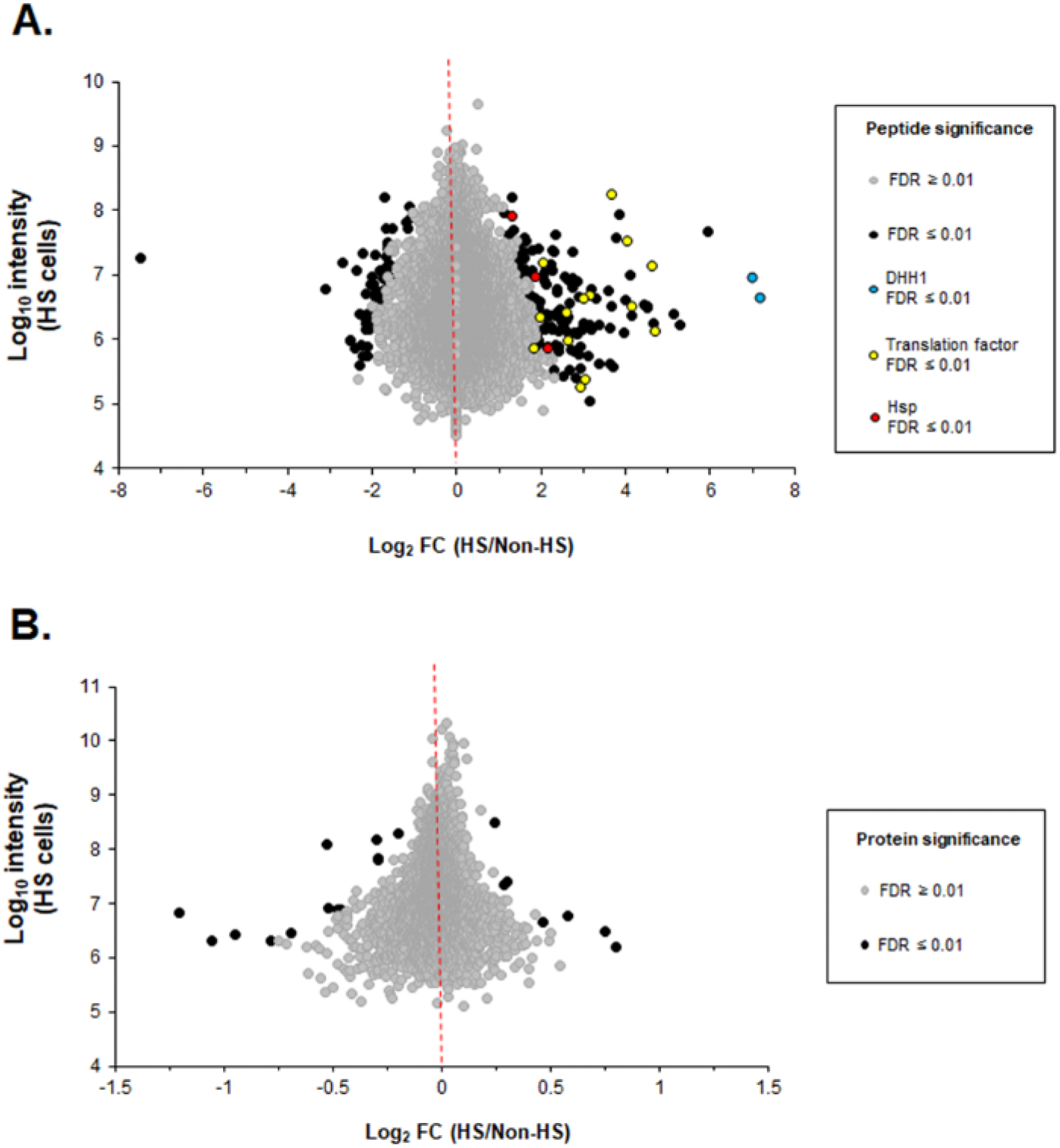
Quantitative proteomic analysis of heat shock response. **A.** Quantification of changes in phosphorylation site abundance. **B.** Quantification of changes in proteins abundance. Plots show change in abundance of phosphorylation sites (HS/untreated) in BSF *T. brucei* cells after 1 h heat shock at 41 °C. Significant changes (FDR ≤ 0.01) following heat shock are indicated as black circles. DHH1 - blue, Translation factors - yellow, Hsps - red.

The 193 HS sensitive phosphorylation sites showed significant categorical enrichment (FDR ≤ 0.01) for 16 curated Gene Ontology (GO) terms (Figure 3). The enriched GO terms include many with anticipated roles in HS response included RNA Helicase activity (FDR 5.9 × 10^-7^), P-body (FDR = 6.8 × 10^-6^), eIF4-F complex (FDR = 6.6 × 10^-6^), nuclear and cytosolic stress granules (FDR = 9.82 × 10^-6^ and 3.5 × 10^-4^) and mRNA binding (FDR = 7.0 × 10^-4^). In addition, there was significant enrichment for known and novel RBPs (FDR 2.7 × 10^-7^) [40, 41], and some enrichment (FDR ≤ 0.05) of HSPs (FDR = 0.013) and Protein kinases (FDR = 0.011) [42, 43].

**Figure 3.**
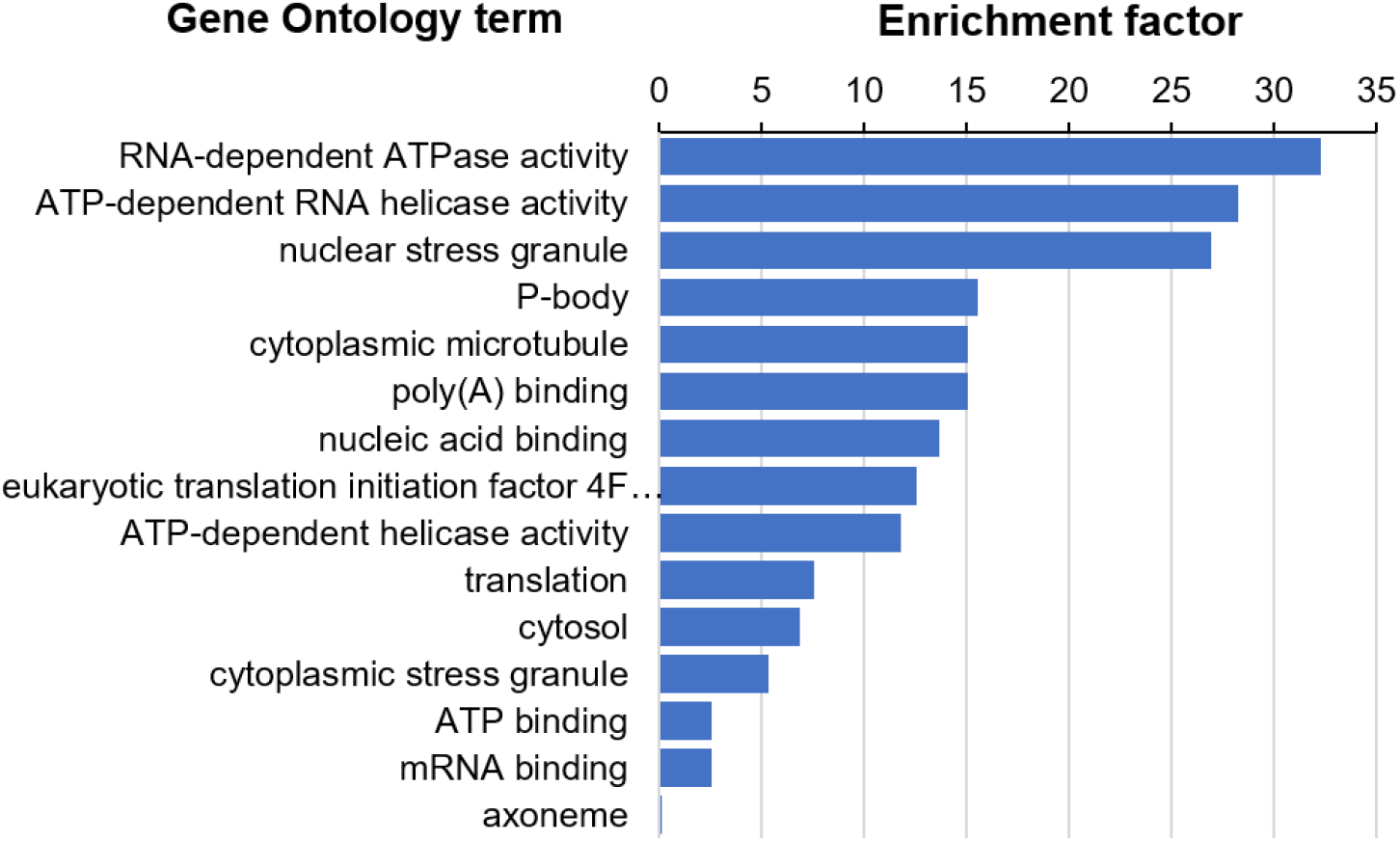
GO term enrichment in heat shock sensitive phosphorylation sites. Enrichment of curated GO terms within the 193 HS sensitive phosphorylation sites was determined using Fisher’s exact test with a Benjamini-Hochberg FDR ≤ 0.01.

Amongst the 20 proteins that changed in abundance in response to HS there was no significant categorical enrichment (FDR ≤ 0.01) of GO terms, and only HSPs were significant enriched (FDR = 0.0078). Interestingly, the abundance of Hsp100 was significantly increased (1.2-fold) in agreement with our qPCR data (≥6-fold, Figure 1C), consistent with a more rapid response occurring at the mRNA level than protein level. These data indicate that protein abundance does not rapidly respond (≤1 h) to heat shock, and that the changes observed are smaller and less widespread than changes in phosphorylation site abundance.

### Phosphorylation modulates the formation of P-bodies and the post-transcriptional regulation of heat shock

The largest increases in phosphorylation site abundance occur on the DEAD-box RNA helicase DHH1 S84 (140-fold) and T82 (126-fold), with an additional site T22 showing a more modest but significant increase (5-fold). In PCF *T. brucei* DHH1 represses certain life cycle stage specific mRNAs via recognition of an AU rich element in the 3’ UTR [44, 45]. Expression of an ATPase deficient DHH1 mutant in PCF increases the number of P-bodies and selectively stabilises certain mRNAs, leading to ISG75 protein accumulation, which we also observe to be significantly increased in abundance upon heat shock (1.4-fold). Upon HS DHH1 is known to re-localise to P-bodies, where it is largely but not entirely co-localised with PAPB2 but not PAPB1 [46]. Phosphorylation of PABP2 was also significantly increased on HS at S93, T167, S212 and S259 by 8- to 16-fold in our data set, whilst phosphorylation of PABP1 was not significantly changed. These data suggest that phosphorylation of DHH1 and PABP2 may play an important role in their re-localisation to P-bodies in response to HS.

DHH1 and PABP2 have been shown to co-immunoprecipitate with the CCCH zinc-finger ZC3H11 (Figure 4), a key regulator of HS response that is essential in BSF but not PCF *T. brucei* [18, 47]. The abundance of ZC3H11 increases after HS, and the protein has been shown to bind to and stabilises the mRNA of many chaperones and Hsps including Hsp70, Hsp83 and Hsp100, in agreement with our qPCR data (figure 1C). A significant increase in phosphorylation of ZC3H11 S275 (16-fold) occurs upon HS, along with three other phosphorylation sites S23, S26 and S279 that were only observed under HS conditions (localisation >0.95). ZC3H11 has previously been reported to be phosphorylated based on changes in protein migration on a gel after phosphatase treatment, but the specific modification sites have not been mapped. We were only able to observe the protein (5 unique peptides) after HS, supporting an increase in abundance, but were unable to quantify the increase in abundance at the protein level. ZC3H11 forms a complex with MKT1, LSM12 and the PABP-binding protein PBP1 [47], but none of these proteins changed significantly at the protein or phosphorylation site level upon HS. MKT1 associates with MAPK6 (NEK6 (Tb927.5.2820) and CK1.2 (Tb927.5.800), and it has been shown that RNAi of CK1.2 reduces the phosphorylation of ZC2H11 under normal and heat shock conditions, whereas RNAi of NEK6 had no effect on ZC3H11 expression [48]. We observe significant increases in phosphorylation of NEK6 S27 (8-fold) and two sites on CK1.2 S19 and S21 (2.8-and 3.5-fold respectively). We also observed significant increases in phosphorylation site abundance on a CCCH zinc-finger protein ZC3H41 that associates with Z3CH11 as well as a DEAD box Helicase (Tb927.3.260), a ubiquitin ligase with a CCCH zinc-finger UPL3 (Tb927.8.1590) and a CCCH zinc-finger proteins ZC3H32 that associate with MKT1 [47] (Figure 4). Taken together, these data suggest that the ZC3H11-MKT1 complex is an important signalling node that is dynamically phosphorylated to modulate RNA binding and P-body formation in response to heat shock.

**Figure 4.**
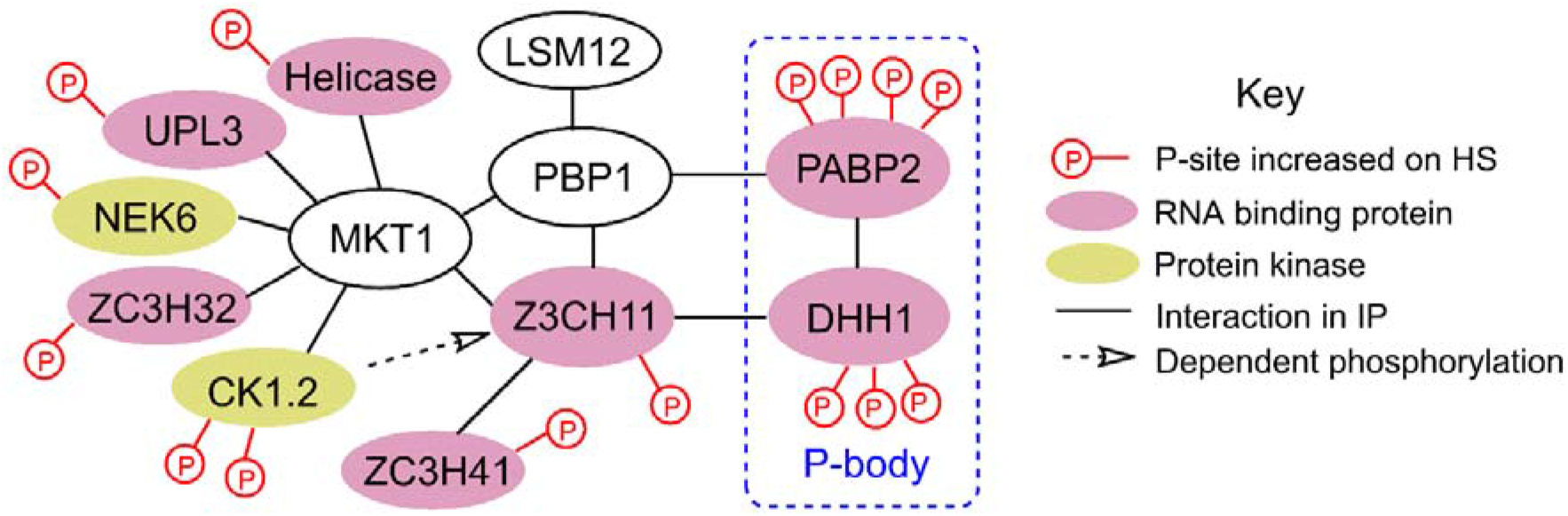
Phosphorylation of the ZC3H11 – MKT1 complex in response to heat shock. Phosphorylation sites that significantly increase in abundance on heat shock (FDR < 0.01) are shown. Interaction data is taken from Singh et al [47].

### Phosphorylation of heat shock proteins occurs more rapidly than changes in abundance

Phosphorylation site on three Hsps were significantly increased after HS: Hsp70.c S464 (Tb927.11.11290, 4.5-fold), Hsp83 S48 (Tb927.11.11960, 3.7-fold) and Hsp70 S79 (Tb927.11.11330, 2.5-fold). None of these three Hsps showed a significant change in protein abundance, but Hsp70 and Hsp83 were shown to have > 2-fold increase at the mRNA level after HS (Figure 1C).

Phosphorylation sites on a further three Hsps (Hsp110, Tb927.10.12710; Hsp84, Tb927.11.250; Hsp83, Tb927.10.10960) were changed > 2-fold upon HS, but were not considered to be significantly changes in our analysis (0.01 > FDR > 0.05), and did not change at the protein level. At the protein level only Hsp100 (Tb927.2.5980) was significantly increased (1.2-fold), in agreement with our qPCR data (Figure 1C). Differential phosphorylation of Hsps, but not changes to mRNA or protein level, is similar to the HS response observed in *Leishmania donovani* [24–26], a kinetoplastid parasite closely related to *T. brucei*.

### Translation factors are phosphorylated in response to heat shock

*We* found significant differential phosphorylation at 18 sites on 9 different proteins that are either eukaryotic translation initiation factors or have an established role in translation (Table 1). The *T.brucei* genome has an expanded complement of components of the eIF4F cap-dependent initiation complex subunits with two eIF4A, six eIF4E and five eIF4G variants, providing an opportunity for greater complexity in translational control. The eukaryotic translation initiation factor 4F complex in *T. brucei* consists of eIF4A-1 – eIF4E-4 – eIF4G-3 which is important for the G1/S transition [49], and a second eIF4A-1 – eIF4E-3 – eIF4G-3/4 complex that may regulate the G2/M transition [33]. HS induced a significant increase in phosphorylation on eIF4A-1 at S51, S101, S147, T155, S237 and T325 by between 3-fold and 24-fold respectively, and a decrease in phosphorylation on eIF4E-4 S90 by 3-fold. Phosphorylation also increases on eIF4G4 S23 4-fold and S68 6-fold, and on Translation associated element 2 S502 (Tb927.11.13060) by 25-fold, proteins shown to interact and localise to stress granules upon starvation [50]. We did not observe any significant change to the phosphorylation of eIF2-a (Tb927.3.2900), consistent with reports that HS acts independently of eIF2α T169 phosphorylation (equivalent to serine 51 in mammalian eIF2α) [8]. Interestingly, the MKT1-like protein MKT1L S1005 (Tb927.10.1490) has the second largest decrease (8.5-fold) in phosphorylation site abundance, and in *Leishmania tarentolae* MKT1L co-purifies with the spliceosome complex responsible for trans-splicing a common spliced leader sequence to all mRNAs [51]. Phosphorylation of proteins involved in translational complexes is suggestive of a mechanism which contributes to the repression of translation and the formation of stress granules/P-bodies in response to HS.

**Table 1.** Heat shock alters the phosphorylation of translational machinery.

| GeneDB | Name | Phosphorylation site | Fold-change |
| --- | --- | --- | --- |
| Tb927.9.4680 | eIF4A-1 | S51<br>S101<br>S147<br>T155<br>S237<br>T325 | 24-fold up<br>17-fold up<br>3.5-fold up<br>3.0-fold up<br>13-fold up<br>7.7-fold up |
| Tb927.6.1870 | eIF4E-4 | S90 | 3.1-fold down |
| Tb927.11.11770 | eIF4E-3 | S424 | 3.5-fold up |
| Tb927.11.10560 | eIF4G-4 | S23<br>S68 | 3.9-fold up<br>6.3-fold up |
| Tb927.8.4500 | eIF4G-5 | S15 | 4.1-fold up |
| Tb927.11.13060 | Translation associated element 2 | S502 | 26.2-fold up |
| Tb927.11.9610 | Translation initiation factor 3i | S248 | 5.8-fold up |
| Tb927.11.2300 | Eukaryotic peptide chain release factor subunit 1 | S172 | 7.1-fold up |
| Tb927.9.10770 | PABP2 | S93<br>T167<br>S212<br>S259 | 16-fold up<br>8.2-fold up<br>8.8-fold up<br>8.1-fold up |

### Phosphorylation of protein kinases in response to heat shock

The *T. brucei* genome contains between 176 – 182 putative protein kinases (PKs) [42, 43], and our data contains 18 phosphorylation sites on 14 PKs that alter significantly in response to heat shock (Table 2), whilst no PKs change significantly at the protein level. PKs with HS sensitive phosphorylation sites include many that have been demonstrated to be essential, and where RNAi ablation in the bloodstream form causes a cell cycle defect, including CK1.2, GSK3, MAPK6, PLK, RCK [52–55]. Amongst the 18 significantly regulated phosphorylation sites, 8 sites decrease in abundance upon HS including the essential AGC S229 (3.2-fold), PLK T198 (3.6-fold) and CDK2 Y57 (3.6-fold). The decrease in phosphorylation of PLK T198 on the T-loop will reduce its activity, preventing progression through the cell cycle as PLK activity is required for cytokinesis. Amongst the 10 sites that show an increase in phosphorylation are GSK3 S44 (4.3-fold) and NEK17 S197 (5.7-fold). Phosphorylation of mammalian GSK3 on S9 reduces its catalytic activity, and it is possible that the S44 phosphorylation adjacent to the N-terminus of the *T. brucei* GSK3 kinase domain has a similar inhibitory role. NEK17 has been identified as being involved in the stumpy induction pathway, with a potential role in cell cycle exit and quiescence [56].

**Table 2.** Phosphorylation of protein kinases in response to heat shock

| GeneDB | Name | Phosphorylation site | Fold change |
| --- | --- | --- | --- |
| Tb927.3.2440 | AGC Essential | S229 | 3.2-fold down |
| Tb927.10.5310 | AMPK | T464 | 3.9-fold down |
|  |  | S468 | 4.0-fold down |
| Tb927.10.3900 | CAMK/CAMKL | S167 | 5.3-fold down |
| Tb927.5.800 | CK1.2 | S21 | 3.5-fold up |
|  |  | S19 | 2.8-fold up |
| Tb927.10.3340 | CMGC/MAPK | T155 | 3.1-fold down |
|  |  | T844 | 4.0-fold down |
| Tb927.7.960 | CMGC/SRPK | S68 | 7.2-fold up |
| Tb927.7.7360 | CRK2 | Y57 | 3.6-fold down |
| Tb927.10.13780 | GSK3 short | S44 | 4.4-fold up |
| Tb927.8.3550 | MAPK3 | Y195 | 4.1-fold up |
| Tb927.10.5930 | NEK17 | S197 | 5.8-fold up |
| Tb927.5.2820 | NEK6 | S27 | 8.1-fold up |
| Tb927.7.6310 | PLK | T198 | 3.6-fold down |
| Tb927.3.690 | RCK | Y177 | 3.0-fold up |
| Tb927.1.1530 | STE group | S731 | 5.7-fold up |

## Discussions

*T. brucei* is an early branching eukaryotic parasite with highly divergent mechanisms for gene regulation compared to its mammalian hosts [22]. As genes with diverse functions are arranged in polycistronic arrays under the transcriptional control of common promoters, *T. brucei* is unable to differentially regulate gene expression at the level of transcription initiation. However, *T. brucei* is subjected to the same physiological pressures as their mammalian hosts. What molecular mechanisms allow for different solutions to common challenges? Our quantitative proteomic and phosphoproteomic analysis shows that during the first hour of HS there is little change in overall protein abundance, but nearly 200 phosphorylation sites are significantly changed in abundance. The differential phosphorylation of thermotolerance proteins in response to HS has previously been reported for the related kinetoplastid *L. donovani* [24–26], but it is unclear how phosphorylation is able to impart thermotolerance.

Our analysis reveals that significantly different phosphorylation in response to HS occurs on RBPs, PKs, translational components, and P-body / stress granule proteins. The decreased phosphorylation of protein kinases with established roles in the cell cycle and increased phosphorylation of translational components such as the eiF4F complex suggests that these phosphorylation sites may play a role in the global translational arrest and exit from the cell cycle that occurs upon HS. The large increase in phosphorylation site abundance on DHH1 and PABP2 may have a role in their sequestration into P-bodies / stress granules in response to heat shock [46]. Both DHH1 and PABP2 have been shown to associate with the ZC3H11-MKT1 complex, along with a number of other RBPs and PKs including CK1.2 that are dynamically phosphorylated in response to HS [47], suggesting the complex may represent a signal integration node.

Following HS, the half-life of both phosphorylated and dephosphorylated ZC3H11 increases, resulting in an increase in the abundance of target mRNAs for Hsps and chaperone proteins [18, 48]. The stability of the dephosphorylated form is greater, and accumulates with prolonged HS. Phosphorylation of ZC3H11 *in vitro* is dependent on the activity of CK1.2, and *in vitro* CK1.2 can phosphorylate N-terminal portions of ZC3H11, but the sites of phosphorylation have not previously been determined [48]. We observed that phosphorylation of ZC3H11 S275 increases 16-fold upon HS, and phosphorylation at S23, S26 and S279 was observed only after HS conditions but we did not observe any sites that occurred only in untreated cells. Of the identified phosphorylation sites, only S23 and S26 occur in the portion of ZC3H11 used in the recombinant kinase assay, and none occur within canonical CK1 recognition motifs, suggesting that the phosphorylation of ZC3H11 by CK1.2 may not be direct. The activity of CK1.2 against ZC3H11 in IP experiments was reduced when cell were treated at 41 °C for 1 hour [48], and we observe an ~3-fold increase in phosphorylation of CK1.2 at S19 and S21 after HS that may be responsible for the decreased activity. Further experiments are required to examine both the temporal dynamic of these phosphorylation sites and to clarify their effect on the HS response.

Temperature changes in the host are a major physiological challenge to parasites and factors conferring HS tolerance constitute overlooked virulence factors. A better understanding of these virulence factors would pave the way for the development of novel drug therapies which selectively target *T. brucei*. This can be achieved either by the development of compounds which directly target Hsps, or via inhibitors of phosphorylation that could specifically target the activation of the HS response.

## Conclusions

Quantitative proteomic and phosphoproteomic analysis of the heat shock response in BSF *T. brucei* has revealed that protein abundance does not rapidly respond (≤1 h) to heat shock, but that significant changes in phosphorylation site abundance occur. We have identified dynamic phosphorylation occurring on RBPs and PKs that may play a role in translational arrest, exit from the cell cycle, P-body formation and upregulation of Hsp and chaperones. The ZC3H11-MKT1 complex appears to represent a key signal integration node in the heat shock response.

## Acknowledgements

The authors would like to acknowledge Calvin Tiengwe for qPCR reagents and discussions. The authors also wish to thank Matthew Child, Paul McKean, Philippe Bastin and Brice Rotureau for helpful discussions. MDU and CB were supported by a BBSRC New Investigator Award (BB/M009556/1) and CPO is employed by the Wellcome Trust (grant no. 095161).

## References

1. Shaw AP, Cecchi G, Wint GR, Mattioli RC, Robinson TP. Mapping the economic benefits to livestock keepers from intervening against bovine trypanosomosis in Eastern Africa. Prev Vet Med 2014; 113(2): 197–210.

2. Simarro PP, Cecchi G, Franco JR, Paone M, Diarra A, Ruiz-Postigo JA, Fevre EM, Mattioli RC, Jannin JG. Estimating and mapping the population at risk of sleeping sickness. PLoS Negl Trop Dis 2012; 6(10):e1859.

3. Baker N, De Koning HP, Maser P, Horn D. Drug resistance in African trypanosomiasis: the melarsoprol and pentamidine story. Trends Parasitol 2013; 29(3): 110–8.

4. Sinha A, Grace C, Alston WK, Westenfeld F, Maguire JH. African trypanosomiasis in two travelers from the United States. Clin Infect Dis 1999; 29(4):840–4.

5. Magona JW, Walubengo J, Olaho-Mukani W, Jonsson NN, Eisler MC. Diagnostic value of rectal temperature of African cattle of variable coat colour infected with trypanosomes and tick-borne infections. Vet Parasitol 2009; 160(3-4):301–5.

6. Franco JR, Simarro PP, Diarra A, Jannin JG. Epidemiology of human African trypanosomiasis. Clin Epidemiol 2014; 6:257–75.

7. Taylor CR, Lyman CP. Heat storage in running antelopes: independence of brain and body temperatures. Am J Physiol 1972; 222(1):114–7.

8. Kramer S, Queiroz R, Ellis L, Webb H, Hoheisel JD, Clayton C, Carrington M. Heat shock causes a decrease in polysomes and the appearance of stress granules in trypanosomes independently of eIF2(alpha) phosphorylation at Thr169. J Cell Sci 2008; 121(Pt 18):3002–14.

9. Muhich ML, Hsu MP, Boothroyd JC. Heat-shock disruption of trans-splicing in trypanosomes: effect on Hsp70, Hsp85 and tubulin mRNA synthesis. Gene 1989; 82(1): 169–75.

10. Chakraborty C, Clayton C. Stress susceptibility in Trypanosoma brucei lacking the RNA-binding protein ZC3H30. PLoS Negl Trop Dis 2018; 12(10):e0006835.

11. Grousl T, Ivanov P, Frydlova I, Vasicova P, Janda F, Vojtova J, Malinska K, Malcova I, Novakova L, Janoskova D, Valasek L, Hasek J. Robust heat shock induces eIF2alpha-phosphorylation-independent assembly of stress granules containing eIF3 and 40S ribosomal subunits in budding yeast, Saccharomyces cerevisiae. J Cell Sci 2009; 122(Pt 12):2078–88.

12. Farny NG, Kedersha NL, Silver PA. Metazoan stress granule assembly is mediated by P-eIF2alpha-dependent and-independent mechanisms. RNA 2009; 15(10):1814–21.

13. Avila CC, Peacock L, Machado FC, Gibson W, Schenkman S, Carrington M, Castilho BA. Phosphorylation of eIF2alpha on Threonine 169 is not required for Trypanosoma brucei cell cycle arrest during differentiation. Mol Biochem Parasitol 2016; 205(1-2):16–21.

14. Chen B, Retzlaff M, Roos T, Frydman J. Cellular strategies of protein quality control. Cold Spring Harb Perspect Biol 2011; 3(8):a004374.

15. Lum R, Tkach JM, Vierling E, Glover JR. Evidence for an unfolding/threading mechanism for protein disaggregation by Saccharomyces cerevisiae Hsp104. J Biol Chem 2004; 279(28):29139–46.

16. Haslberger T, Zdanowicz A, Brand I, Kirstein J, Turgay K, Mogk A, Bukau B. Protein disaggregation by the AAA+ chaperone ClpB involves partial threading of looped polypeptide segments. Nat Struct Mol Biol 2008; 15(6):641–50.

17. Wendler P, Shorter J, Snead D, Plisson C, Clare DK, Lindquist S, Saibil HR. Motor mechanism for protein threading through Hsp104. Mol Cell 2009; 34(1):81–92.

18. Droll D, Minia I, Fadda A, Singh A, Stewart M, Queiroz R, Clayton C. Post-transcriptional regulation of the trypanosome heat shock response by a zinc finger protein. PLoS Pathog 2013; 9(4):e1003286.

19. Costa-Martins AG, Lima L, Alves JMP, Serrano MG, Buck GA, Camargo EP, Teixeira MMG. Genome-wide identification of evolutionarily conserved Small Heat-Shock and eight other proteins bearing alpha-crystallin domain-like in kinetoplastid protists. PLoS One 2018; 13(10):e0206012.

20. Hombach A, Ommen G, Macdonald A, Clos J. A small heat shock protein is essential for thermotolerance and intracellular survival of Leishmania donovani. J Cell Sci 2014; 127(Pt 21):4762–73.

21. Mahat DB, Salamanca HH, Duarte FM, Danko CG, Lis JT. Mammalian Heat Shock Response and Mechanisms Underlying Its Genome-wide Transcriptional Regulation. Mol Cell 2016; 62(1):63–78.

22. Clayton C. The regulation of trypanosome gene expression by RNA-binding proteins. PLoS Pathog 2013; 9(11):e1003680.

23. Singh A, Minia I, Droll D, Fadda A, Clayton C, Erben E. Trypanosome MKT1 and the RNA-binding protein ZC3H11: interactions and potential roles in post-transcriptional regulatory networks. Nucleic Acids Res 2014; 42(7):4652–68.

24. Morales MA, Watanabe R, Dacher M, Chafey P, Osorio Y, Fortea J, Scott DA, Beverley SM, Ommen G, Clos J, Hem S, Lenormand P, Rousselle JC, Namane A, Spath GF. Phosphoproteome dynamics reveal heat-shock protein complexes specific to the Leishmania donovani infectious stage. Proc Natl Acad Sci U S A 2010; 107(18):8381–6.

25. Tsigankov P, Gherardini PF, Helmer-Citterich M, Spath GF, Zilberstein D. Phosphoproteomic analysis of differentiating Leishmania parasites reveals a unique stage-specific phosphorylation motif. J Proteome Res 2013; 12(7):3405–12.

26. Tsigankov P, Gherardini PF, Helmer-Citterich M, Spath GF, Myler PJ, Zilberstein D. Regulation dynamics of Leishmania differentiation: deconvoluting signals and identifying phosphorylation trends. Mol Cell Proteomics 2014; 13(7): 1787–99.

27. Minia I, Merce C, Terrao M, Clayton C. Translation Regulation and RNA Granule Formation after Heat Shock of Procyclic Form Trypanosoma brucei: Many Heat-Induced mRNAs Are also Increased during Differentiation to Mammalian-Infective Forms. PLoS Negl Trop Dis 2016; 10(9):e0004982.

28. Livak KJ, Schmittgen TD. Analysis of relative gene expression data using real-time quantitative PCR and the 2(-Delta Delta C(T)) Method. Methods 2001; 25(4):402–8.

29. Domingo-Sananes MR, Szoor B, Ferguson MA, Urbaniak MD, Matthews KR. Molecular control of irreversible bistability during trypanosome developmental commitment. J Cell Biol 2015; 211(2):455–68.

30. Urbaniak MD, Martin DM, Ferguson MA. Global quantitative SILAC phosphoproteomics reveals differential phosphorylation is widespread between the procyclic and bloodstream form lifecycle stages of *Trypanosoma brucei*. J Proteome Res 2013; 12(5):2233–44.

31. Wisniewski JR, Zougman A, Nagaraj N, Mann M. Universal sample preparation method for proteome analysis. Nat Methods 2009; 6(5):359–62.

32. Ruprecht B, Koch H, Medard G, Mundt M, Kuster B, Lemeer S. Comprehensive and reproducible phosphopeptide enrichment using iron immobilized metal ion affinity chromatography (Fe-IMAC) columns. Mol Cell Proteomics 2015; 14(1):205–15.

33. Benz C, Urbaniak MD. Organising the cell cycle in the absence of transcriptional control: Dynamic phosphorylation co-ordinates the *Trypanosoma brucei* cell cycle post-transcriptionally. PLoS Pathog 2019; 15(12):e1008129.

34. Schroeder MJ, Shabanowitz J, Schwartz JC, Hunt DF, Coon JJ. A neutral loss activation method for improved phosphopeptide sequence analysis by quadrupole ion trap mass spectrometry. Anal Chem 2004; 76(13):3590–8.

35. Cox J, Mann M. MaxQuant enables high peptide identification rates, individualized p.p.b.-range mass accuracies and proteome-wide protein quantification. Nat Biotechnol 2008; 26(12):1367–72.

36. Cox J, Neuhauser N, Michalski A, Scheltema RA, Olsen JV, Mann M. Andromeda: a peptide search engine integrated into the MaxQuant environment. J Proteome Res 2011; 10(4):1794–805.

37. Aslett M, Aurrecoechea C, Berriman M, Brestelli J, Brunk BP, Carrington M, Depledge DP, Fischer S, Gajria B, Gao X, Gardner MJ, Gingle A, Grant G, Harb OS, Heiges M, Hertz-Fowler C, Houston R, Innamorato F, Iodice J, Kissinger JC, Kraemer E, Li W, Logan FJ, Miller JA, Mitra S, Myler PJ, Nayak V, Pennington C, Phan I, Pinney DF, Ramasamy G, Rogers MB, Roos DS, Ross C, Sivam D, Smith DF, Srinivasamoorthy G, Stoeckert CJ, Jr., Subramanian S, Thibodeau R, Tivey A, Treatman C, Velarde G, Wang H. TriTrypDB: a functional genomic resource for the Trypanosomatidae. Nucleic Acids Res 2010; 38(Database issue):D457–62.

38. Vizcaino JA, Cote RG, Csordas A, Dianes JA, Fabregat A, Foster JM, Griss J, Alpi E, Birim M, Contell J, O’kelly G, Schoenegger A, Ovelleiro D, Perez-Riverol Y, Reisinger F, Rios D, Wang R, Hermjakob H. The PRoteomics IDEntifications (PRIDE) database and associated tools: status in 2013. Nucleic Acids Res 2013; 41(Database issue):D1063–9.

39. Tyanova S, Temu T, Sinitcyn P, Carlson A, Hein MY, Geiger T, Mann M, Cox J. The Perseus computational platform for comprehensive analysis of (prote)omics data. Nat Methods 2016; 13(9):731–40.

40. Erben ED, Fadda A, Lueong S, Hoheisel JD, Clayton C. A genome-wide tethering screen reveals novel potential post-transcriptional regulators in Trypanosoma brucei. PLoS Pathog 2014; 10(6):e1004178.

41. Lueong S, Merce C, Fischer B, Hoheisel JD, Erben ED. Gene expression regulatory networks in Trypanosoma brucei: insights into the role of the mRNA-binding proteome. Mol Microbiol 2016; 100(3):457–71.

42. Parsons M, Worthey EA, Ward PN, Mottram JC. Comparative analysis of the kinomes of three pathogenic trypanosomatids: Leishmania major, Trypanosoma brucei and Trypanosoma cruzi. BMC Genomics 2005; 6:127.

43. Urbaniak MD, Mathieson T, Bantscheff M, Eberhard D, Grimaldi R, Miranda-Saavedra D, Wyatt P, Ferguson MA, Frearson J, Drewes G. Chemical proteomic analysis reveals the drugability of the kinome of Trypanosoma brucei. ACS Chem Biol 2012; 7(11):1858–65.

44. Kramer S, Queiroz R, Ellis L, Hoheisel JD, Clayton C, Carrington M. The RNA helicase DHH1 is central to the correct expression of many developmentally regulated mRNAs in trypanosomes. J Cell Sci 2010; 123(Pt 5):699–711.

45. Kramer S, Carrington M. An AU-rich instability element in the 3’UTR mediates an increase in mRNA stability in response to expression of a dhh1 ATPase mutant. Translation (Austin) 2014; 2(1):e28587.

46. Kramer S, Bannerman-Chukualim B, Ellis L, Boulden EA, Kelly S, Field MC, Carrington M. Differential localization of the two T. brucei poly(A) binding proteins to the nucleus and RNP granules suggests binding to distinct mRNA pools. PLoS One 2013; 8(1):e54004.

47. Singh A, Minia I, Droll D, Fadda A, Clayton C, Erben E. Trypanosome MKT1 and the RNA-binding protein ZC3H11: interactions and potential roles in post-transcriptional regulatory networks. Nucleic Acids Research 2014; 42(7):4652–4668.

48. Minia I, Clayton C. Regulating a Post-Transcriptional Regulator: Protein Phosphorylation, Degradation and Translational Blockage in Control of the Trypanosome Stress-Response RNA-Binding Protein ZC3H11. PLoS Pathog 2016; 12(3):e1005514.

49. An T, Liu Y, Gourguechon S, Wang CC, Li Z. CDK Phosphorylation of Translation Initiation Factors Couples Protein Translation with Cell-Cycle Transition. Cell Rep 2018; 25(11):3204–3214 e5.

50. Fritz M, Vanselow J, Sauer N, Lamer S, Goos C, Siegel TN, Subota I, Schlosser A, Carrington M, Kramer S. Novel insights into RNP granules by employing the trypanosome’s microtubule skeleton as a molecular sieve. Nucleic Acids Research 2015; 43(16):8013–8032.

51. Tkacz ID, Gupta SK, Volkov V, Romano M, Haham T, Tulinski P, Lebenthal I, Michaeli S. Analysis of spliceosomal proteins in Trypanosomatids reveals novel functions in mRNA processing. J Biol Chem 2010; 285(36):27982–99.

52. Urbaniak MD. Casein kinase 1 isoform 2 is essential for bloodstream form Trypanosoma brucei. Mol Biochem Parasitol 2009; 166(2): 183–5.

53. Jones NG, Thomas EB, Brown E, Dickens NJ, Hammarton TC, Mottram JC. Regulators of Trypanosoma brucei cell cycle progression and differentiation identified using a kinome-wide RNAi screen. PLoS Pathog 2014; 10(1):e1003886.

54. Hammarton TC, Kramer S, Tetley L, Boshart M, Mottram JC. Trypanosoma brucei Polo-like kinase is essential for basal body duplication, kDNA segregation and cytokinesis. Mol Microbiol 2007; 65(5):1229–48.

55. Wei Y, Li Z. Distinct roles of a mitogen-activated protein kinase in cytokinesis between different life cycle forms of Trypanosoma brucei. Eukaryot Cell 2014; 13(1): 110–8.

56. Mony BM, Macgregor P, Ivens A, Rojas F, Cowton A, Young J, Horn D, Matthews K. Genome-wide dissection of the quorum sensing signalling pathway in Trypanosoma brucei. Nature 2014; 505(7485):681–5.

57. Ridewood S, Ooi CP, Hall B, Trenaman A, Wand NV, Sioutas G, Scherwitzl I, Rudenko G. The role of genomic location and flanking 3’UTR in the generation of functional levels of variant surface glycoprotein in *Trypanosoma brucei*. Mol Microbiol 2017; 106(4):614–634.

