## Supplementary data for "Phosphoproteomic analysis of mammalian infective *Trypanosoma brucei* subjected to heat shock suggests atypical mechanisms for thermotolerance"

**Supplementary material**

*Content:*

Table S1. qPCR primers for *T. brucei Hsps.*

Figure S1. Ribosomal RNA integrity is maintained in BSF *T. brucei* for up to 2 h after heat shock.

*Separately available:*

Table S2. Phosphoproteomic analysis of heat shock response in BSF *T. brucei*. (.xlsx)

Table S3. Proteomic analysis of heat shock response in BSF *T. brucei*. (.xlsx)

Table S1. qPCR primers for *T. brucei Hsps.*

| **Target transcript** | **Forward primer** | **Reverse primer** | **Previous publication** |
| --- | --- | --- | --- |
| *rDNA 28s * | GTAAGTTCGCAAGAAGCAT | ACCAGAAGGAGGTTAGTAGATA | [7] |
| *Hsp70* (Tb927.11.11330) | GCGGGGACGATTGCTGGTCT | GCACGTTGCGTTCCTTCCCC |  |
| *Hsp83* (Tb927.10.10980) | AGGCCGGTGGCGATATGAGC | AACGGTTACGCGGTCTGCGA |  |
| *Hsp110* (Tb927.10.12710) | TCCACGTTGCGGGAGAAGCA | GGTCGCTTCACAACCGCACG |  |
| *Hsp40* (Tb927.10.8540) | GCGTTTCGTTGGCGAGGGTG | GAACCGCGGATGCGGCTTTT |  |
| *Hsp100* (Tb927.2.5980) | ATGCGGGTACGGTGCTGGAG | GGGCCTCTGCTCCGGAAGTG |  |

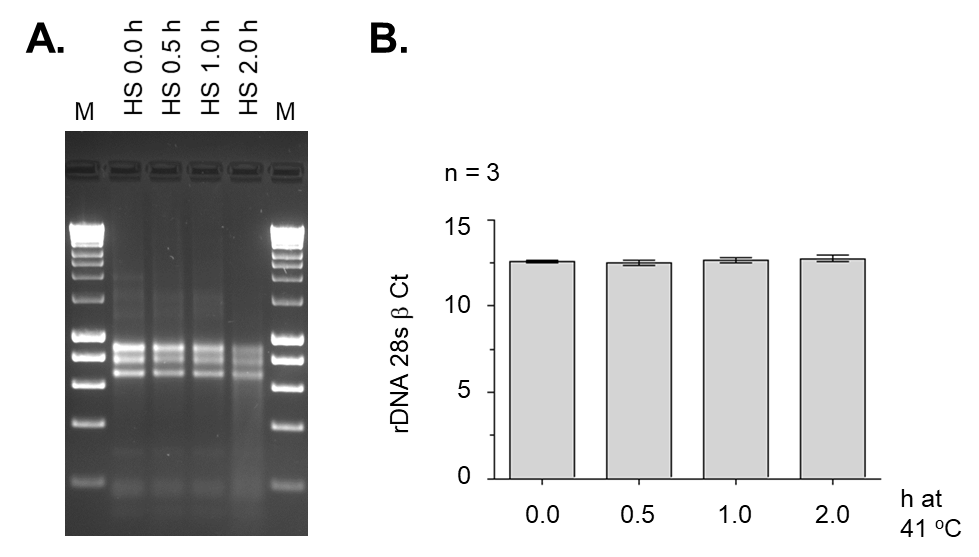

Figure S1. Ribosomal RNA integrity is maintained in BSF *T. brucei* for up to 2 h after heat shock. A. 300 ng of total RNA from heat shocked (HS) *T. brucei* visualised on a 2 % agarose gel. B. The Ct (critical threshold) values for the rDNA 28s β subunit measured using qPCR. Error bars are of standard deviation from three biological replicates.
